# OpEnHiMR: Optimization based Ensemble Model for Prediction of Histone Modification in Rice

**DOI:** 10.1101/2025.11.10.687509

**Authors:** Dipro Sinha, Sneha Murmu, Abhik Sarkar, Md Yeasin, Sougata Bhattacharjee, Himanshushekhar Chaurasia, Dwijesh Chandra Mishra, Neeraj Budhlakoti, Sudhir Srivastava, Sunil Archak, Aparna Kumari, Girish Kumar Jha

**Affiliations:** ICAR-Indian Agricultural Statistics Research Institute, New Delhi, Delhi-110012; The Graduate School, ICAR-Indian Agricultural Research Institute, New Delhi, Delhi-110012; ICAR-National Institute for Plant Biotechnology, New Delhi, Delhi-110012; ICAR-Indian Agricultural Research Institute, Hazaribagh, Jharkhand-825405; ICAR-Central Institute for Research on Cotton Technology, Mumbai – 400019; ICAR-National Bureau of Plant Genetic Resources, New Delhi, Delhi-110012; Centre for Cellular and Molecular Biology, Hyderabad-500007

**Keywords:** Epigenetics, Histone modification, Methylation, Acetylation, Ensemble model

## Abstract

Histone modifications are central to gene regulation, yet their systematic identification in plants remains limited due to the complexity of epigenomic landscapes. We present OpEnHiMR, an optimization-based ensemble learning framework for multiclass prediction of three key histone modifications, H3K4me3, H3K27me3, and H3K9ac, in rice. The framework integrates Support Vector Machines, Random Forest, and Gradient Boosting models, optimized via Ant Colony Optimization to maximize performance. Biologically meaningful features, including mononucleotide binary encoding, nucleotide chemical properties, GC content, and k-mer frequencies, were used for training after rigorous data curation and redundancy removal. OpEnHiMR achieved a classification accuracy of 77.54%, outperforming individual models and ensuring improved recall, specificity, and Matthews correlation coefficient. Model interpretability was enhanced using SHAP analysis, which highlighted critical sequence features influencing prediction outcomes. To promote community-wide adoption, a user-friendly webserver (https://dipro-sinha.shinyapps.io/OpEnHiMR/) and R package (https://cran.r-project.org/web/packages/OpEnHiMR/index.html) were developed. OpEnHiMR thus provides a scalable, accurate, and interpretable tool for histone modification prediction in plants, advancing epigenomics research and supporting data-driven crop improvement strategies.

## Introduction

The comprehensive knowledge of epigenetic modifications in plants, encompassing DNA methylation, RNA methylation, and histone modifications [1] in regulating gene expression, is not completely ingrained. However, it is well understood that chromatin architecture tightly regulates DNA methylation and histone modifications, thereby selectively activating or repressing gene expression [2]. In DNA methylation, methylations at the fourth and fifth positions on the pyrimidine ring of cytosine [4-methylcytosine (4mC) and 5-methylcytosine (5mC)], as well as the sixth position on the adenine purine ring [N6-adenine methylation or N6-methyladenine (6mA)], rank among the most prevalent DNA modifications [3], [4] [(Yuan, 2020); (Zhang et al., 2007)] Nevertheless, the investigation of 6mA has been limited owing to its sporadic distribution throughout genomes. The precise whereabouts and operational mechanism of 6mA in eukaryotes are primarily unknown. On the other hand, histones are basic proteins involved in DNA damage repair, replication, recombination, and essential functions such as gene expression regulation [5], [6]. In addition to acetylation, methylation, and phosphorylation, histones can be covalently modified in various ways. The histone code hypothesis posits that unique covalent modifications occurring on one or more tails of histones operate sequentially or in combination to generate a “histone code”, which is interpreted by other proteins to initiate specific downstream events [7]. True epigenetic codes capable of inheritance may consist of histone codes, which may be impermanent or more stable in nature[8]. It is noteworthy that histone deacetylation and histone H3 lysine 27 trimethylation (H3K27me3) play a role in repressing transcription in eukaryotes [9]. In contrast, histone acetylation and H3K4me3 have been inevitably linked to the stimulation of gene expression, which significantly influences plant development and plays a role in plant responses to biotic and abiotic stresses. Again, H3K9me is prevalently linked to transcriptional repression, whereas H3K36me and H3K9 acetylation serve as an indicator of gene activation [10]. Many established experimental techniques are employed to quantify the abundance of different epigenetic marks on the genome-wide scale. For instance, chromatin immunoprecipitation sequencing (ChIP-seq) for histone modifications and DNA bisulfite sequencing (BS-seq) for 5mC and 6mA, respectively, and single-molecule real-time sequencing (SMRT-seq) and methylated RNA immunoprecipitation sequencing (MeRIP-seq) are a few examples [11]. Research employing these techniques has unveiled novel insights into the interplay between transcriptional regulation and epigenetics, illuminating patterns of variation showing epigenetic diversity in a population, thereby inaugurating a ‘big-data era of epigenetics [12]’. However, epigenomic modifications differ from genomics research in that they are dynamic in nature. Therefore, the scope of experiments is typically restricted to particular tissues, genotypes, or environmental conditions, restricting the depth of investigation. Utilising *in silico* methodologies to examine potential sites of epigenomic modifications in plants is a logical course of action.

Recent advancements in computational biology have led to the development of various tools aimed at predicting histone modifications, as summarized in Table 1. These tools utilize diverse machine learning (ML) and deep learning (DL) approaches, incorporating genomic sequences, chromatin accessibility data, and other epigenetic features to identify histone modification sites[13]. However, most of these tools are limited in their applicability to multiclass prediction, generalizability across species, and interpretability in plant systems. Tools such as DeepHistone [14], iHMnBS [15], SMEP (Smart Model for Epigenetics in Plants) [16] (Y. Wang et al., 2021), Histone-Net [17], dHICA [11] and KAS Former [18] and have demonstrated significant potential in histone modification prediction. DeepHistone, for example, integrates DNA sequences and chromatin accessibility to achieve high accuracy, while Histone-Net introduces advanced sequence representation methods for histone marker prediction. However, these tools are constrained by high computational demands, limited multiclass prediction capabilities, and optimization challenges specific to plant systems. Furthermore, their focus has primarily been on human and mammalian datasets, leaving plant histone modifications relatively underexplored.

**Table 1:** Comparative Review of Related Tools.

| Tool/<br>Method | Features | ML/DL<br>Techniques | Advantages | Disadvantages | Performance | References |
| --- | --- | --- | --- | --- | --- | --- |
| DeepHistone | Sequence, chromatin accessibility | Convolutional Neural Networks | High accuracy, interpretable features | Limited to specific dataset | ROC-AUC: ~0.94 | Yin et al., 2019 |
| Histone-Net | k-mer embeddings, DNA2Vec representations | Multi-label classification | Multi-task prediction, cross-domain applicability | High dimensionality | Accuracy: 0.69 | Asim et al., 2023 |
| SMEP (Smart Model for Epigenetics in Plants) | Sequential feature by One Hot Encoding | Convolutional Neural Network (CNN) | Multiple types of epigenetic modification predictions (DNA methylation, RNA methylation and Histone modification) | Insufficient for accurately characterizing the entire complement of epigenomic modifications occurring at a genome-wide scale | AUC: H3K4me3: 0.94; H3K27me3: 0.93; H3K9ac: 0.96 | Wang et al., 2021 |
| iHMnBS | Local and global features extracted from input DNA sequences | Convolutional Neural Network and variants of Recurrent Neural Networks (RNNs) | Can identify 7 types of histone modifications that need not be anchored in the central sequence exclusively, Can predicts the binding | Lack of natural language processing approaches (NLP), Lack of information about the DNA and histone | AuROC: About 0.95 (Acc. To cell lines), Average AuROC (acc. To histone | Lyu et al., 2022 |
|  |  |  | sites of DNA double helix structures to modified histones | binding site-specific Histone modification | modifications): 0.94 |  |
| dHICA | DNA Sequence data, Chromatin Accessibility | Convolutional Neural Network (CNN) along with Transformer block | Multiple Histone Modification prediction at Different cell lines, capture long-range interactions between genomic elements | Noise and variability in input data due to lack of data transformation or normalization | Average AuPRC; 0.73 | Wang *et al.*, 2024 |
| KASFormer | DNA Sequential feature by One Hot Encoding, KAS Seq signals | Convolutional Neural Network (CNN) and Transformer block | Able to capture long range interactions between genomic elements, Cross cell and cross species prediction capabilities | Trained on only single cell line | AUC: H3K36me3: 0.8525; H4k20me1: 0.7850 | Jia *et al.*, 2024 |

**Table 2:** Details of parameters and packages used for developing the best three ML models.

| Machine Learning Model | Parameters | R-Package |
| --- | --- | --- |
| Support Vector Machine | SVM-Type = C-classification, SVM-Kernel = radial | e1071 |
| Random Forest | Number of trees: 500, split: 5 | randomForest |
| Gradient Boosting | Method = Stochastic, n.trees = 150, interaction.depth = 3, shrinkage = 0.1 and n.minobsinnode = 10 | gbm |

**Table 3:** Performance Evaluation of the Models (Randomization: Linear; K=1)

| Model | Accuracy | Specificity | Recall | MCC |
| --- | --- | --- | --- | --- |
| SVM | 0.7398 | 0.8519 | 0.6277 | 0.4806 |
| Regression Tree | 0.6596 | 0.7997 | 0.5194 | 0.3205 |
| Random Forest | 0.7412 | 0.8508 | 0.6316 | 0.4825 |
| Naive Bayes | 0.6169 | 0.775 | 0.4588 | 0.2288 |
| Multiple Logistic Reg. | 0.716 | 0.8388 | 0.5932 | 0.4345 |
| kNN | 0.7044 | 0.8264 | 0.5825 | 0.4086 |
| Gradient Boosting | 0.7427 | 0.8533 | 0.6321 | 0.4866 |
| ANN | 0.6244 | 0.775 | 0.4739 | 0.2547 |

In this study, we harnessed the power of machine learning to address the challenge of predicting epigenetic modifications in plants, focusing on three critical histone modification classes: H3K4me3 (associated with gene activation) [19], H3K27me3 (linked to transcriptional repression) [20], and H3K9ac (a marker of active transcription) [21]. By leveraging advanced computational approaches, we developed the first multiclass prediction tool specifically tailored for plant epigenomics. This represents a significant breakthrough, as most existing tools are designed for binary classification or are optimized for non-plant species.

To evaluate the predictive capabilities of various algorithms, we systematically tested multiple machine learning models. Among these, Support Vector Machines (SVM) [22], Random Forest (RF) [23], and Gradient Boosting (GB) [24] demonstrated superior performance in distinguishing the three classes of histone modifications. These models excelled in terms of accuracy, specificity, recall, and F1-score across diverse datasets, affirming their robustness in addressing the multiclass nature of this prediction task.

Building on these findings, we further enhanced predictive performance through the development of an. This bio-inspired optimization approach effectively combined the strengths of the top-Ant Colony Optimization (ACO)-based ensemble model [25] performing individual classifiers, fine-tuning their contributions to maximize accuracy. Our ensemble model achieved an impressive accuracy of 77.54%, significantly surpassing the standalone models and demonstrating its potential for addressing complex classification problems in epigenomics.

The ability of our tool to predict multiple histone modification classes simultaneously offers a comprehensive framework for investigating the intricate regulatory roles of H3K4me3, H3K27me3, and H3K9ac in plants. These modifications are pivotal in shaping gene expression, influencing plant development, and mediating responses to environmental stresses. By providing a multiclass prediction capability, our tool opens new avenues for exploring the dynamic interplay of histone modifications in plants [26], a domain that remains underexplored compared to other epigenomic studies. Furthermore, our approach integrates experimental insights with in silico methodologies, addressing the inherent limitations of traditional experimental techniques, such as their dependency on specific tissues, genotypes, or environmental conditions. The computational efficiency and scalability of our tool make it a valuable resource for analysing large-scale epigenomic datasets, facilitating deeper insights into plant biology. This study represents a significant advancement in plant epigenomics, bridging the gap between experimental and computational research.

## Materials and Methods

### Data Collection

The data for this study was curated to build a robust foundation for training and evaluating our machine learning models. Positive data, representing validated instances of histone modifications H3K4me3, H3K27me3, and H3K9ac, was sourced from the SMEP database [16] (https://www.elabcaas.cn/smep/index.html). This database contains experimentally verified histone modification sites, ensuring a reliable source for constructing the positive dataset.

To generate the negative dataset, we used sequence randomization techniques [29] which involved creating artificial sequences that retained certain statistical properties of the original sequences but were not associated with histone modifications. Three distinct methods were employed for randomization:

1. Linear Randomization: Sequences were shuffled randomly without considering any dependencies between adjacent nucleotides.
2. Markov Chain Randomization: A first-order Markov model was used to preserve the immediate dependencies between adjacent nucleotides, ensuring a more realistic nucleotide composition.
3. Euler Path-Based Randomization: A Eulerian graph approach was applied to retain higher-order k-mer frequencies, preserving local sequence patterns more effectively.

For each randomization method, k-mer lengths (k) ranging from 1 to 3 were used, leading to the generation of 9 distinct types of negative datasets (3 randomization methods × 3 k-mer settings). This comprehensive strategy ensured that the negative dataset was both diverse and biologically realistic, minimizing potential biases that could arise from using a single randomization method. By combining experimentally validated positive data with systematically generated negative datasets, we created a balanced and biologically relevant training and testing dataset. This dataset is designed to effectively train machine learning models to distinguish true histone modification sites from randomized non-modified sequences, ensuring robust model performance.

### Data Preprocessing

Effective data preprocessing is an essential step in building reliable machine learning models, particularly when analysing epigenetic datasets. In this study, we employed a combination of redundancy removal and data balancing to ensure the quality and representativeness of our dataset. To remove redundant sequences and reduce dataset complexity, we utilized CD-HIT (Cluster Database at High Identity with Tolerance) [30]. CD-HIT clusters sequences based on sequence similarity, retaining only representative sequences within each cluster. For our analysis, we set the sequence identity threshold to 60%, which ensures that sequences with more than 60% similarity are clustered, and only one representative sequence from each cluster is retained. This lower threshold was chosen to prioritize sequence diversity while removing redundant or highly similar sequences that could introduce bias into the model. The redundancy removal was applied independently to each class of histone modifications (H3K4me3, H3K27me3, H3K9ac) and the negative dataset. This ensured that all datasets were processed in a consistent manner, resulting in a non-redundant dataset that preserved biological diversity. Following redundancy removal, it was important to address the class imbalance in the dataset to avoid biased model training. A random sampling approach was used to balance the number of sequences in each class (Table S2 – S4).

Additionally, sequence padding was performed to ensure that all sequences had the same length. Padding involves adding dummy characters (typically a placeholder like “N”) to sequences that are shorter than the maximum sequence length, making them uniform in size. This step is critical for ensuring that the input data is compatible with machine learning algorithms, which typically require fixed-length inputs.

### Feature Extraction

After removing the redundant data, the non-redundant dataset is initialized for feature extraction. Mononucleotide Binary Encoding (MBE) [31] has been considered as an essential feature for this study due to its relevance to histone modification lies in its ability to efficiently encode sequence features that are critical for identifying sequence motifs associated with histone-modifying enzyme binding (e.g., CpG dinucleotides near H3K27me3 marks). MBE represents DNA sequences as a string of binary digits [32]. These nucleotides are adenine (A), guanine (G), cytosine (C), and thymine (T). In mononucleotide binary encoding, each nucleotide is assigned a four-bit code where “A” is represented as 1000, “C” as 0010, “G” as 0100, and “T” as 0001.

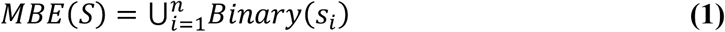

Where *S* = {*s*_1_, *s*_2_, …, *s*_*n*_} is the DNA sequence, *s*_*i*_,is the ith nucleotide in the sequence and *Binary*(*s*_*i*_) *maps the nucleotide*(*s*_*i*_) *to its corresponding binary value*

Apart from MBE, another significant feature for nucleotide sequences is Nucleotide Chemical Properties (NCP). NCP [33] lies on the fact that nucleotide bases have different chemical characteristics. These characteristics can be represented through three properties, *viz*. ring structure, hydrogen bond, and amino/keto bases. Each property is being given a value of 0 or 1 depending on its two alternative possibilities. A, C, G, and T have been represented in this study by the ring structure, hydrogen bond, and amino/keto bases, which were (1, 1, 1), (0, 0, 1), (1, 0, 0), and (0, 1, 0), respectively.

GC content is one of the frequently used sequential features in this context. The ratio of the abundance of G and C bases with respect to sequence length is known as GC content. GC content of a sequence can be represented as

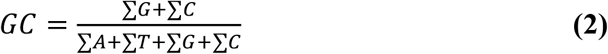

As the GC content values are greater than other features it results in relatively decreased algorithmic efficacy. As a solution, the GC content value has been normalised in this study by taking the log of the GC values with base as *e*. This log transformed feature is known as Log GC content (*GC*).

In case of prediction of histone modification oligo nucleotide frequency plays a pivotal role because it reflects underlying patterns in DNA sequence that affect chromatin structure, transcription factor binding as well as the recruitment of histone-modifying enzymes. Among all oligonucleotides frequency Di Nucleotide Frequency (DNF) and Tri Nucleotide Frequency (TNF) [34]are most important ones in the context of histone modification. DNF and TNF can be calculated as per the formula [35] (Goldman, 1993) as mentioned below in the equations (4) and (5).

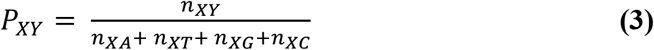

where *n*_*XY*_ = No. of times the dinucleotide XY is observed in the sequence, *P*_*XY*_ = Di Nucleotide Frequency, (n_*XA*_ + *n*_*XT*_ + *n*_*XG*_ + *n*_*XC*_) **=** Total no. of possible dinucleotides.

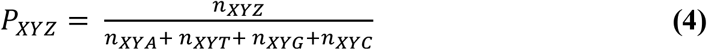

where *n*_*XYZ*_ = No. of times the trinucleotide XYZ is observed in the sequence, *P*_*XYZ*_ = Tri Nucleotide Frequency, (*n*_*XYA*_ + *n*_*XYT*_ + *n*_*XYG*_ + *n*_*XYC*_) **=** Total no. of possible trinucleotides.

### Feature Selection

These five most influential features have been extracted from the dataset and fed to two distinguished feature selection criterions based on 1) Random Forest and 2) Stepwise regression algorithm.

Random Forest-based feature selection [36] is a technique used to identify the most important features for predictive modelling by leveraging the power of Random Forest algorithms. For feature selection, Random Forest evaluates the importance of each feature based on how it contributes to the prediction accuracy across all trees. One common method is calculating the Mean Decrease in Impurity (MDI), which measures how much each feature reduces the Gini impurity (5) in the decision-making process. The features that consistently lead to higher reductions in impurity are considered more important.

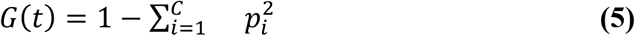

where C is the number of classes, and p_i_ is the proportion of class i at node t. Then the total importance I(f) for a feature f is given in equation (6)

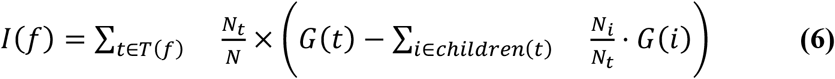

where, Nt is the number of samples at node t, N is the total number of samples, children(t) are the child nodes of t and G(i) is the Gini impurity of child node i.

Another method is Stepwise regression [37] which is a popular method for feature selection in statistical modelling. It involves iteratively adding or removing predictors based on their statistical significance to build a model that best explains the relationship between the dependent and independent variables. Basic formula for multiple linear regression algorithm is mentioned in the equation 7.

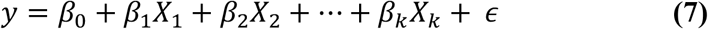

where y is the dependent (response) variable, *X*_1_, *X*_2_, *X*_3_, ……., *X*_*k*_ are the predictor (independent) variables included in the model, *β*_0_ is the intercept, *β*_1_, *β*_2_, ……, *β*_*k*_ are the coefficients of the selected features and ϵ is the error term. The primary objective of this algorithm is to identify the most relevant variables while avoiding overfitting or underfitting.

After obtaining selected features from both of above-mentioned feature selection criterion separately an ensemble module has been implemented. In this ensemble technique only those features which have been selected by both of the criterions *i*.*e. Features*_*RF*_ and *Features*_*SWR*_ are considered as finally selected features for model development **(8)**.

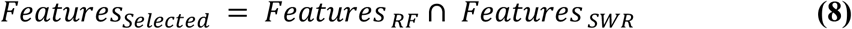

### Model Building

Nine machine learning models were considered for this multiclass classification research [38], [39], [40], namely Support Vector Machine (SVM) [22], Decision Tree (DT) [41], Random Forest (RF) [23], k-Nearest Neighbour (k-NN) [42], Artificial Neural Network (ANN) [43], AdaBoost [44], Naïve Bayes[45], Multiple Linear Regression (MLR) [46], and Gradient Boosting (GB) [24]. Each model has its own advantages and limitations in addressing classification tasks. Our objective was to identify the models that performed best for the multiclass histone modification dataset.

To validate model performance, 10-fold cross-validation (CV) was performed on the entire dataset. The results of cross-validation provided an additional layer of robustness in selecting the best-performing models.

After performance evaluations, the top three models, based on accuracy, were selected for further analysis. These were SVM, RF, and GB, which were then used to construct the final ensemble model. A brief description of these three models is provided below:

#### Support Vector Machine

Support Vector Machine (SVM) [22] is a versatile and robust machine learning method widely used for classification tasks, especially with high-dimensional biological data. In this research, SVM effectively handled multiclass classification by transforming data into a high-dimensional feature space and constructing an optimal hyperplane for separating classes. A one-vs-one approach was used for multiclass handling. For implementation, we used the “e1071” package in the R programming language with a linear kernel.

Grid search was applied to tune the hyperparameters of the SVM model. The grid search process explored different combinations of kernel functions and cost parameters. The model was evaluated based on accuracy, and the hyperparameters yielding the highest accuracy were selected for further analysis [47].

#### Random Forest

Random Forest (RF) [23]is another widely recognized machine learning method for high-dimensional and complex biological datasets. It combines adaptive nearest neighbours with bagging to improve prediction accuracy and stability. RF constructs an ensemble of decision trees, and for multiclass classification, it uses a majority voting mechanism to assign a class to each sequence. This method effectively handles correlated features and class imbalances in the data. The RF model was implemented in the R environment using the “randomForest” package. For Random Forest, grid search [47] was conducted to find the optimal values for the number of trees, maximum tree depth, and other relevant parameters. The grid search evaluated the models based on accuracy, and the best-performing set of hyperparameters was selected.

#### Gradient Boosting

Gradient Boosting (GB) [24]is a sequential machine learning algorithm designed to improve model accuracy by iteratively adding weak learners and minimizing the prediction error. For multiclass classification, GB uses a log-loss function to optimize performance across all classes. The GB model was implemented using the “gbm” package in R.

Grid search was used to fine-tune the hyperparameters [47] of all the models, including the number of trees, learning rate, and tree depth. The search evaluated performance based on accuracy, and the hyperparameter set with the highest accuracy was selected for further analysis. To ensure optimal performance for multiclass classification, hyperparameter tuning using grid search was performed for all three models (SVM, RF, and GB). The hyperparameters were adjusted to achieve the highest accuracy on the multiclass histone modification dataset. The final set of tuned hyperparameters used for training these models is provided in Table 5.

### Ensemble Learning

In this research, Ant Colony Optimization (ACO) (Shi et al., 2011) was employed to optimize the weight parameters for an ensemble model aimed at improving the classification performance for a multiclass dataset. The objective was to combine the outputs of four different classes using weighted coefficients and maximize the accuracy of the ensemble model. The following steps outline the approach used, including the details of the optimization process.

The ensemble model is constructed by combining the outputs of four classifiers, each corresponding to a different class. The objective function (Obj) computes the weighted sum of features from the dataset for each class, based on a set of weights, and then calculates the accuracy of the predicted class using these weights.

For each solution (set of weights), the following steps are performed:

1. The weighted sum for each class (*Cls*_1_, *Cls*_2_, *Cls*_3_, *Cls*_4_) is computed by applying the weights *w*[1] and *w*[2] to the corresponding columns in the dataset.
2. The class with the highest probability is predicted for each sample using which.max function.
3. The accuracy (ACC) of the predicted classes is calculated by comparing the predicted labels with the actual labels from the dataset (Data1[,1]).

The goal is to optimize the weight values *w*[1] and *w*[2] to maximize this accuracy.

The ACO algorithm is used to optimize the weights of the ensemble model to maximize accuracy. The key steps of ACO are as follows:

1. Initialize pheromone levels: *pheromone* [*i*] = 0.1 *for each parameter i*
2. Initialize *best_solution* = *NULL*
3. Initialize *best_score* = *NULL*
4. For *iter* = 1 *to max_iter* do:
  a. For *ant* = 1 *to n_ants* do:
    i. For *param* = 1 *to n_params* do:
      1. Generate random value for pheromone-based weight:

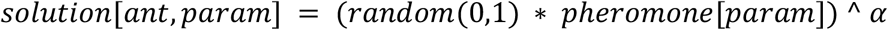
      2. Normalize solution such that *sum(solution[ant*,]) <= 1
    ii. Evaluate the solution using the objective function:

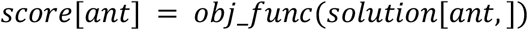
  b. Find best solution in this iteration:

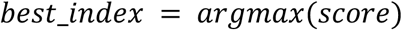 If *score*[*best_index*] > *best_score* then:

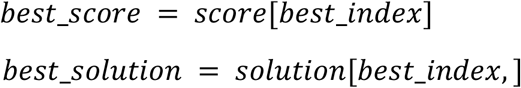
  c. Update pheromone levels for each parameter:

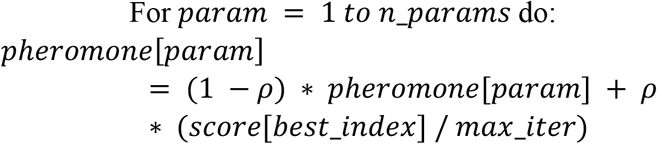
5. Return *best_solution* and *bect_score*

### Performance Evaluation

In the other experiments, sensitivity, specificity, accuracy, and MCC have been used as evaluation metrics for the classifiers [48], [49], [50], [51], [52], [53], [54], [55], [56], the same have been considered in this manuscript.

The proportion of positively tagged cases that are projected to be positive is the sensitivity.

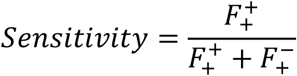

The proportion of negatively tagged cases that are projected to be negative is the sensitivity.

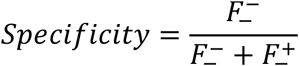

The ratio of successfully identified cases to all test data points is known as the accuracy.

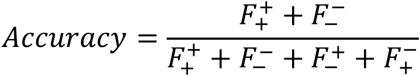

Between the actual and predicted series, there is a correlation known as the Matthews correlation coefficient (MCC). It returns numbers between −1 and +1. A value of 0 is similar to a random forecast, while a coefficient of −1 signifies a full difference between the prediction and the observation. A coefficient of +1 denotes a flawless prediction. The MCC can be calculated directly from the confusion matrix by the formula:

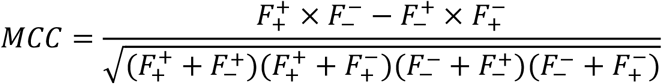

where,

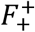 = Instances that are true predicted as true.

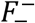 = Instances that are false predicted as false.

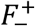 = Instances that are false predicted as true.

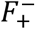 = Instances that are true predicted as false.

## Results

### Feature Selection Outcomes

The feature selection process was carried out in two distinct stages, utilizing two different methods: RF and SWR. First, the RF-based feature selection module was employed to assess the importance of each feature in predicting the target variable. In this approach, the importance of features was determined by calculating the Mean Decrease in Impurity (MDI) [27]score, which reflects how much each feature contributes to reducing the Gini impurity across all decision trees in the forest. Features that led to significant reductions in impurity were considered more important. This allowed us to rank features based on their relevance to the model.

Simultaneously, a Stepwise Regression approach was applied to the dataset. SWR works by iteratively adding or removing features based on their statistical significance in explaining the relationship between the dependent variable and the predictors. This method ensures that only the most statistically relevant features are retained, helping to avoid overfitting or underfitting in the model. Through this iterative process, SWR selects a subset of features that contribute the most to the model’s explanatory power.

Once the features were selected by both RF and SWR individually, the next step was to take the intersection of these two sets of features (Figure 1). This step ensures that only the features that were selected by both methods—indicating their importance and statistical relevance—are considered for further analysis. From this intersection, the top 50 most important features were selected, as they were consistently identified as relevant by both RF and SWR. This final set of top 50 features was then used for the development of the predictive model, ensuring that only the most reliable and informative features, validated by two different selection techniques, were included in the analysis. This combined approach enhances the robustness of the feature selection process and provides a more reliable set of features for model training.

**Figure 1:**
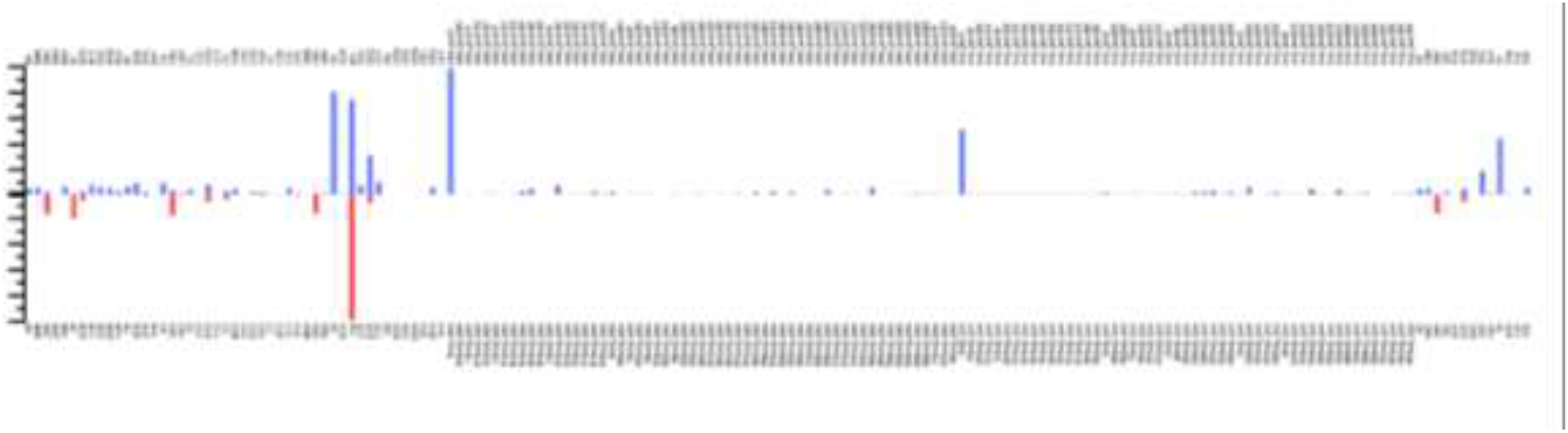
Importance score of features obtained from SwR and RF.

### Probabilistic-feature analysis

We employed t-SNE [28] with the following parameters: perplexity = 50, n_dims = 3, and learning rate = 500 to visualize the data before and after feature selection for both the training and testing sets. The perplexity value of 50 was chosen to balance the local and global aspects of the data, helping to capture both the local neighbourhood relationships and the broader global structure in the lower-dimensional space. The n_dims = 3 setting allowed us to visualize the data in three dimensions, providing a more intuitive understanding of how the data points are distributed. A learning rate of 500 was applied to ensure faster convergence of the algorithm and to avoid the risk of getting stuck in local minima. By applying t-SNE before and after feature selection, we were able to observe the impact of the feature selection process on the data’s clustering and separability, helping us identify how well the retained features facilitated the differentiation between classes in both the training and testing sets (Figure 2).

**Figure 2:**
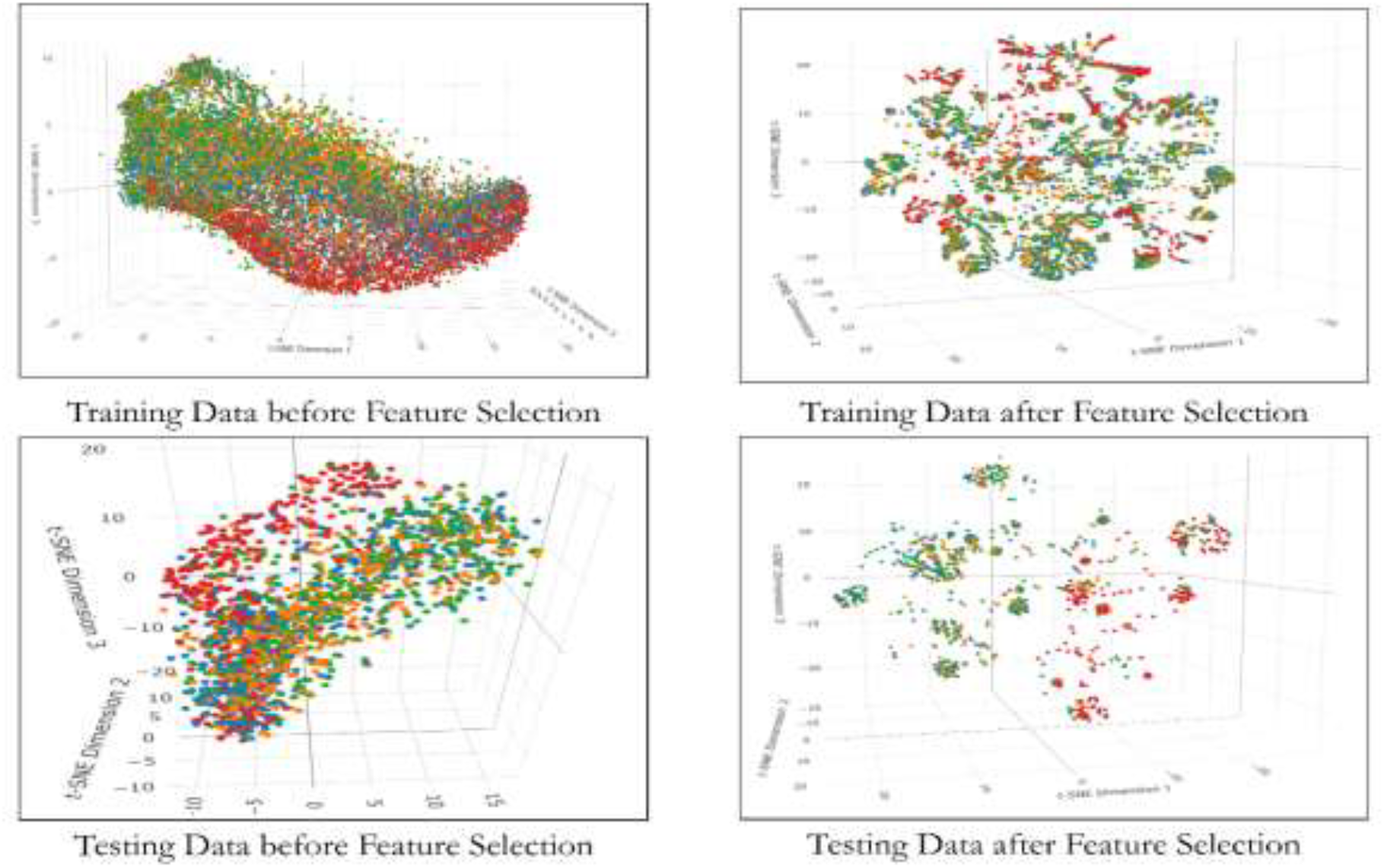
Impact of Feature Selection on Training and Testing Data Visualization.

### Model Performance

Each model’s performance is evaluated using standard metrics such as accuracy, specificity, recall, and MCC. The best results were observed with linear randomization with K value of 1 (Table S6). The performance of the models on nine datasets are given in the supplemental tables (Table S1-S9).

The table compares the performance of eight machine learning models across key metrics: accuracy, specificity, recall, and MCC. Random Forest, Gradient Boosting, and SVM emerge as the top performers, with high accuracy (around 74%) and specificity (85%), demonstrating strong overall classification capabilities, particularly in minimizing false positives and maximizing both true positives and negatives. These models also show high recall, effectively identifying positive instances. Naive Bayes and ANN, however, exhibit lower accuracy, specificity, recall, and MCC, indicating they are less reliable, particularly in detecting positive cases and maintaining a balance between true and false classifications. Multiple Logistic Regression and kNN show moderate performance, with kNN excelling in recall but lagging in MCC, while Multiple Logistic Regression maintains a balanced performance. Overall, Random Forest and Gradient Boosting are the most reliable choices, suitable for tasks that require both high precision and recall, whereas Naive Bayes and ANN are less effective for this dataset.

### Model Interpretation

We employed SHAP (SHapley Additive exPlanations) (Nohara et al., 2019) to interpret the predictions of our machine learning models and to identify the most influential features contributing to classification performance. SHAP assigns a contribution value (SHAP value) to each feature, quantifying its impact on the model’s output while ensuring a fair distribution of importance across all features by considering feature interactions. As illustrated in the SHAP summary plots (Figure 3), certain features consistently exhibited strong contributions across models. Notably, ‘mbe_pos1’ and ‘ncp_pos1’ emerged as the most influential features in all three models—Gradient Boosting, Random Forest, and SVM demonstrating a significant and consistent impact on predictions. Additionally, several k-mer patterns, such as ‘CC’, ‘TA’, ‘AT’, and ‘AAA’ also played notable roles, although their relative influence varied between models.

**Figure 3:**
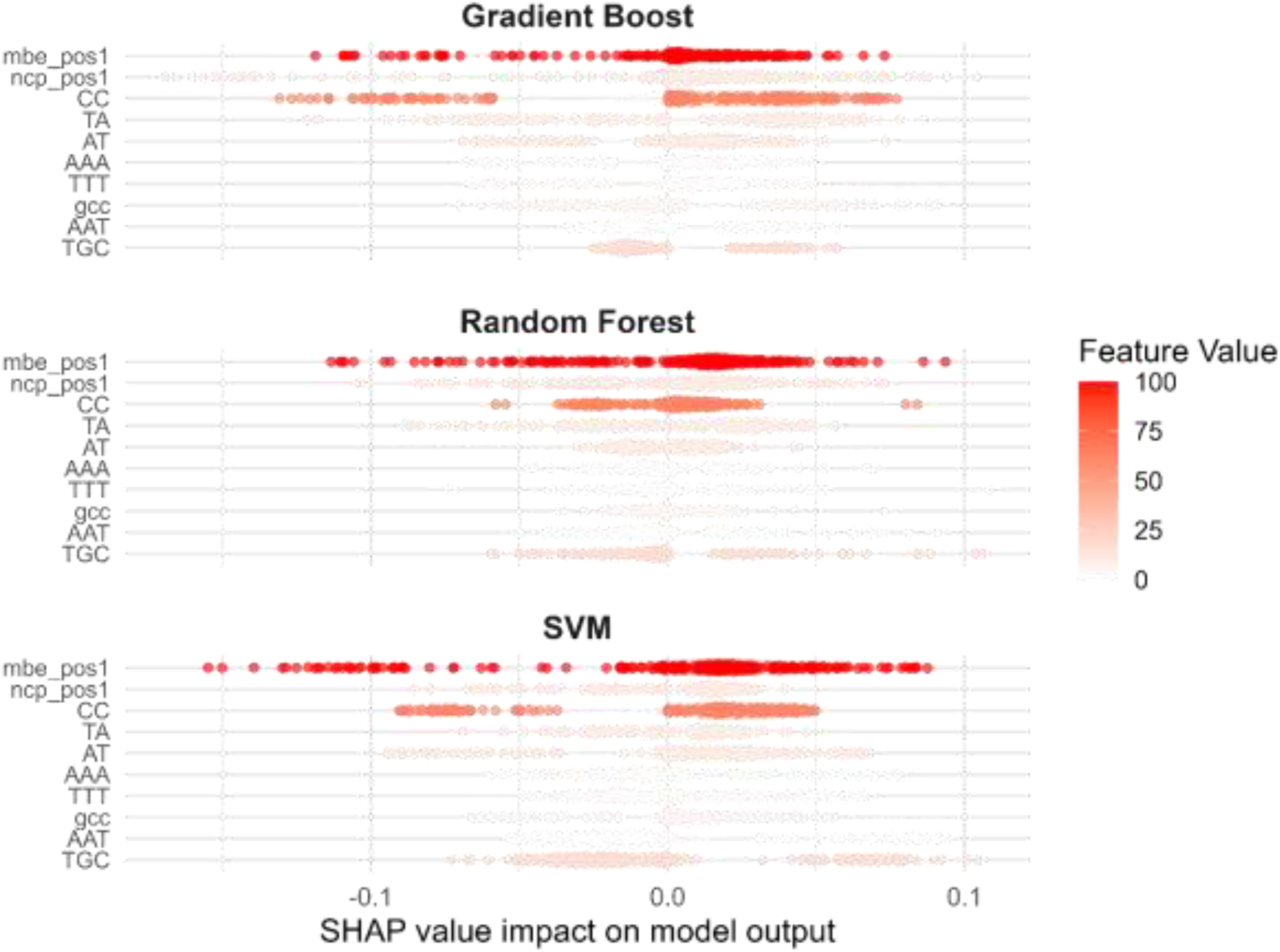
SHAP values for features for OpEnHiMR - Colors indicate feature values (gradient of red), and SHAP (positive and negative) values indicate the directionality of the features.

The plot also reveals the directionality and distribution of feature effects. For instance, higher values of ‘mbe_pos1’ were associated with a positive SHAP contribution, suggesting a strong association with the predicted class. The similarity in SHAP value distributions across models indicates robustness in feature relevance. Overall, SHAP facilitated a transparent understanding of model behaviour, enabling both qualitative insights and model refinement by highlighting dominant and potentially redundant features. These insights were further instrumental in evaluating model generalizability and minimizing potential biases.

### Performance Evaluation of OpEnHiMR

OpEnHiMR achieved the overall accuracy of 0.7754, demonstrating good classification performance. This suggests that the model has little bias towards either class and is successful in differentiating between positive and negative cases. Its dependability is further supported by the Matthews Correlation Coefficient (MCC), which offers a fair assessment of classification performance at 0.5534 (Figure 4). This implies that OpEnHiMR ensures reliable and strong predictions by maintaining a good trade-off between true positives, true negatives, false positives, and false negatives. Furthermore, this MCC score indicates superior predictive consistency compared to Random Forest (0.5465) and SVM (0.5317). In applications where identifying genuine positives is critical, the model’s recall of 0.6585 demonstrates its competence in identifying positive cases. More thorough identification of the target condition is ensured by a higher recall value, which results in fewer positive cases being overlooked. This recall value further supports OpEnHiMR’s better capacity to catch positive occurrences, outperforming both Random Forest (0.6541) and SVM (0.6387). Furthermore, OpEnHiMR’s specificity of 0.8924 indicates that it can accurately detect negative cases, hence lowering false positives. This is particularly helpful in situations like fraud detection or medical diagnostics where the expense of false positives is considerable. OpEnHiMR exhibits a minor but significant boost in specificity when compared to Random Forest (0.8902) and Gradient Boosting (0.8879), guaranteeing more precise identification of negative cases.

**Figure 4:**
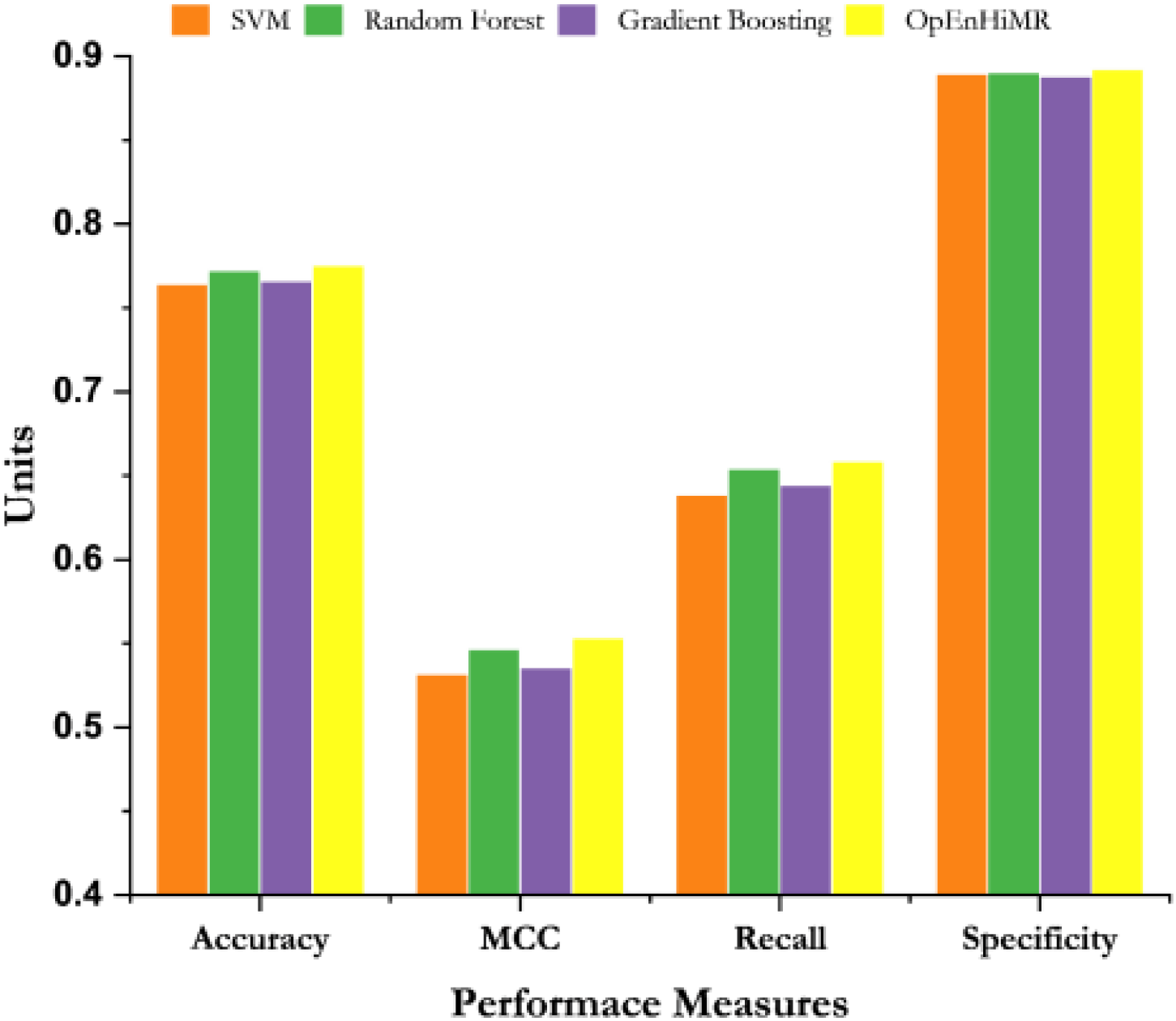
Performance Comparison of OpEnHiMR with the Top Three Models.

### Tool Development

We developed a user-friendly webserver (https://dipro-sinha.shinyapps.io/OpEnHiMR/), shown in Figure 5, and an associated R package (https://cran.r-project.org/web/packages/OpEnHiMR/index.html) to make our machine learning models and data analysis tools more accessible to a broader audience. The webserver offers an intuitive interface that allows users to interact with the models directly without needing to delve into the underlying code or complex configurations. It supports various functionalities such as data uploading, model training, prediction, and visualization, streamlining the workflow for both experienced data scientists and those less familiar with the technical aspects of machine learning.

**Figure 5:**
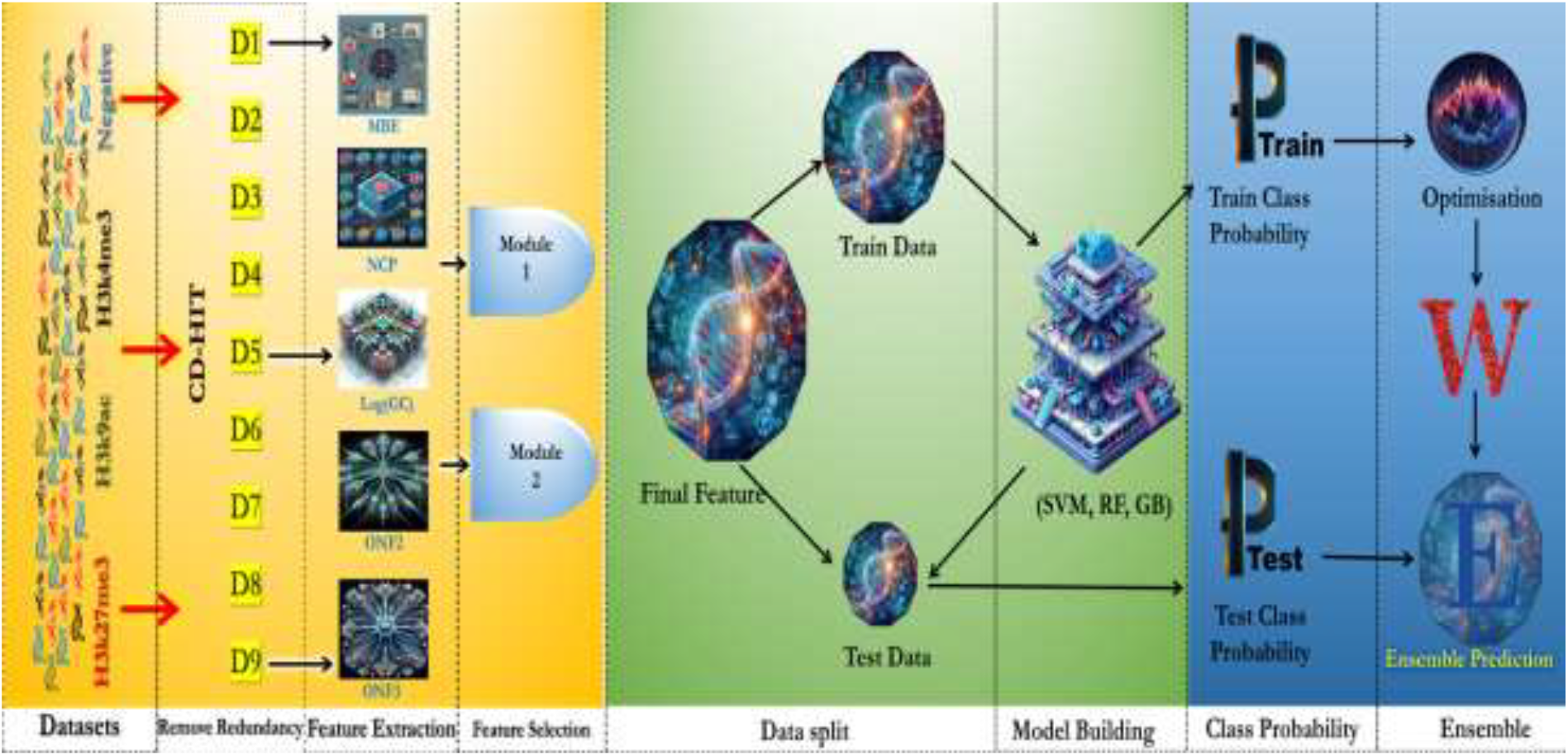
Overall Workflow of OpEnHiMR.

The accompanying R package, which has garnered around 7100 downloads, provides a more programmatic approach for users who prefer working with R. The package includes a comprehensive set of functions for data manipulation, model training, feature selection, and interpretation using SHAP values, among other features. Its high download count reflects its utility and adoption by the bioinformatics and broader research community. By combining the ease of the webserver with the flexibility of the R package, we’ve ensured that our tools are both accessible and versatile, supporting a wide range of use cases from exploratory data analysis to complex model development and interpretation.

## Discussion

The study presents a significant advancement in the field of epigenomics, particularly in understanding histone modifications through a novel machine learning approach. This research addresses the complexities associated with predicting histone modifications in rice, utilizing an ensemble model that integrates various machine learning techniques. The findings highlight several key aspects that contribute to the broader understanding of epigenetic regulation in plants.

The use of ensemble learning, combining Support Vector Machines (SVM), Random Forest (RF), and Gradient Boosting (GB), demonstrates a robust methodology for improving predictive accuracy in classifying histone modifications. The ensemble model, enhanced by Ant Colony Optimization (ACO), not only outperformed individual classifiers but also provided a comprehensive framework for multiclass prediction. This approach is particularly noteworthy as it allows for simultaneous prediction of multiple histone modification classes, which is crucial for understanding their regulatory roles in gene expression. Histone modifications play a pivotal role in regulating gene expression and influencing plant development. The study emphasizes the importance of specific modifications—H3K4me3, H3K27me3, and H3K9ac—in mediating responses to environmental stresses and developmental cues. By effectively predicting these modifications, the proposed tool can facilitate deeper insights into how plants adapt to changing environments, which is increasingly relevant in the context of climate change and agricultural sustainability. The rigorous data collection and preprocessing methods employed in this study enhance the reliability of the results. The combination of experimentally validated positive data with systematically generated negative datasets ensures that the machine learning models are trained on a balanced and biologically relevant dataset. Furthermore, the redundancy removal and class balancing techniques employed help mitigate biases that could compromise model performance. This research represents a significant step forward in computational epigenomics by bridging experimental and computational methodologies. The ability to predict histone modifications using an *in-silico* approach addresses the limitations of traditional experimental techniques, which often require specific tissues or conditions for analysis. The scalability and computational efficiency of the developed tool make it a valuable resource for analysing large-scale epigenomic datasets, thereby opening new avenues for research. The implications of this study extend beyond basic research into potential applications in personalized medicine and crop improvement strategies. By providing insights into the dynamic interplay of histone modifications, this tool can aid in developing strategies for enhancing plant resilience and productivity. Future research could focus on expanding the model to include additional epigenetic marks or applying it to other plant species, further enriching our understanding of plant epigenomics.

## Conclusion

This study introduces OpEnHiMR, a robust and interpretable machine learning framework tailored for the multiclass prediction of histone modifications in rice. By integrating biologically relevant sequence-based features with advanced ensemble learning techniques optimized through Ant Colony Optimization, the model successfully predicts three key histone marks associated with gene activation and repression. OpEnHiMR consistently outperforms standalone models in accuracy, recall, specificity, and MCC, reflecting its effectiveness in addressing the complexity of plant epigenomic datasets. The incorporation of SHAP-based interpretability further enhances the biological relevance and transparency of the model. Through the development of a publicly accessible webserver and R package, OpEnHiMR democratizes access to powerful epigenomic prediction tools, enabling broader applications in plant functional genomics. This work lays the foundation for future expansion into additional epigenetic marks and plant species, ultimately contributing to precision breeding and the understanding of epigenetic regulation in crops.

## Data Availability

All the datasets used and codes developed to conduct this experiment are available at https://github.com/snehaiasri/OpEnHiMR/

## Acknowledgements

The authors gratefully acknowledge the CABin grant (F. no. Agril. Edn.4-1/2013-A & P), Indian Council of Agricultural Research, Ministry of Agriculture and Farmers’ Welfare, Govt. of India, and ICAR-Indian Agricultural Statistics Research Institute, New Delhi, for providing facilities and support. The authors also acknowledge the Department of Biotechnology, Govt. of India, for the BIC project grant (BT/PR40161/BTIS/137/32/2021).

## Author Contributions

D.S., D.C.M., S.A. and G.K.J. conceived and designed the experiments; D.S., S.M., M.Y., H.C., S.B., and A.S. performed the experiments; D.S., S.M., D.C.M., N.B., H.C., and S.S. analysed the data; D.S., A.S., S.B., M.Y., S.A. and G.K.J. wrote the paper. All authors discussed the results and commented on the manuscript.

## Supplemental Information

**Table S1:** Performance Evaluation of Dataset 1 (Randomization: Euler; K= 1)

| Model | Accuracy | Specificity | Recall | MCC |
| --- | --- | --- | --- | --- |
| SVM | 0.7146 | 0.8300 | 0.5992 | 0.4285 |
| Regression Tree | 0.6231 | 0.7597 | 0.4865 | 0.2457 |
| Random Forest | 0.7195 | 0.8307 | 0.6083 | 0.4371 |
| Naive Bayes | 0.6502 | 0.7762 | 0.5242 | 0.2992 |
| Multiple Logistic Regr. | 0.6583 | 0.7939 | 0.5226 | 0.3158 |
| kNN | 0.6813 | 0.7981 | 0.5646 | 0.3565 |
| Gradient Boosting | 0.7170 | 0.8322 | 0.6018 | 0.4337 |
| ANN | 0.5991 | 0.7801 | 0.4180 | 0.2630 |

(The Table S1 describes the comparative analysis of performance of different machine learning models for Dataset 1 with Randomization: Euler; K= 1)

**Table S2:**
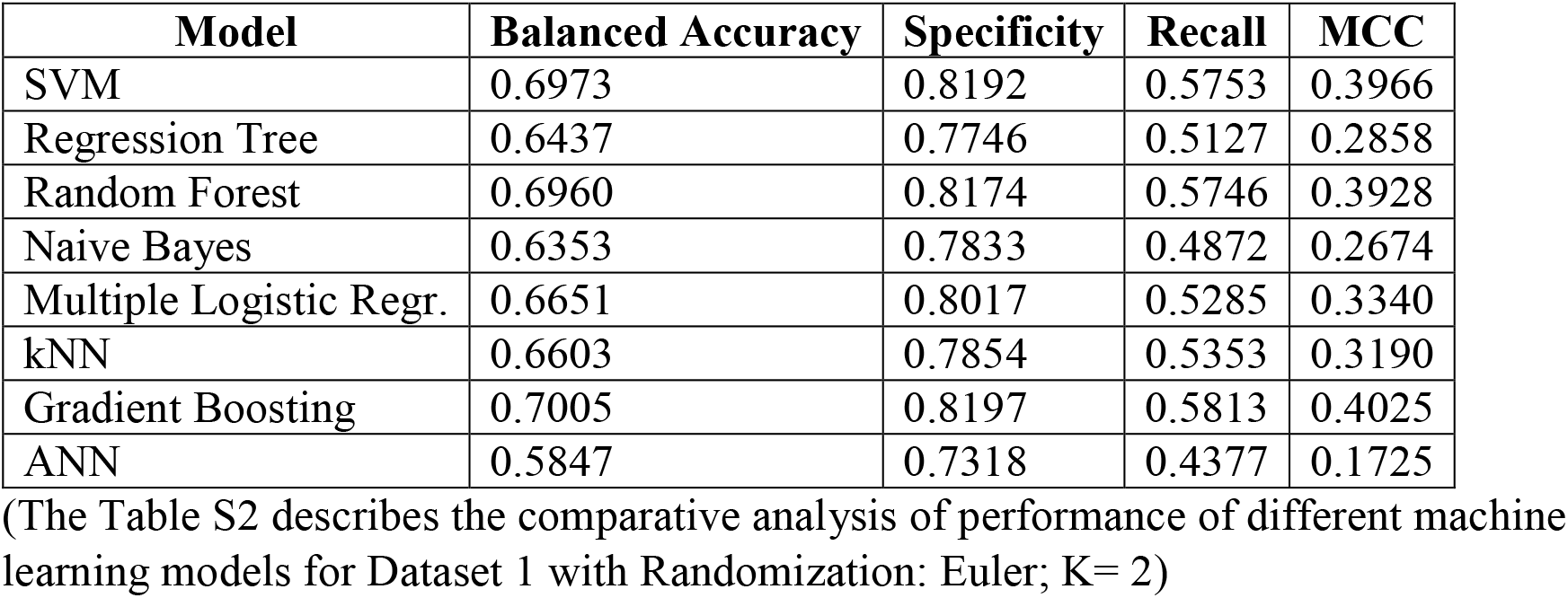
Performance Evaluation of Dataset 1 (Randomization: Euler; K= 2)

**Table S3:**
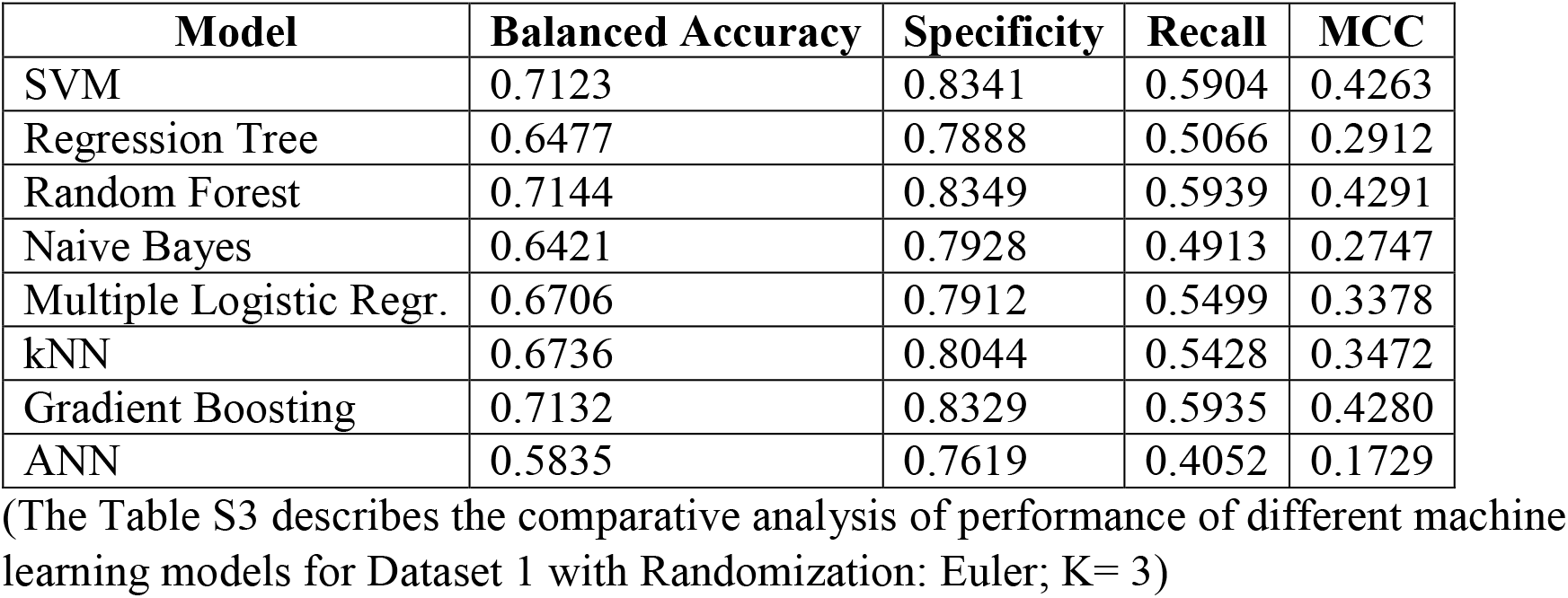
Performance Evaluation of Dataset 1 (Randomization: Euler; K= 3)

| Model | Balanced Accuracy | Specificity | Recall | MCC |
| --- | --- | --- | --- | --- |
| SVM | 0.7123 | 0.8341 | 0.5904 | 0.4263 |
| Regression Tree | 0.6477 | 0.7888 | 0.5066 | 0.2912 |
| Random Forest | 0.7144 | 0.8349 | 0.5939 | 0.4291 |
| Naive Bayes | 0.6421 | 0.7928 | 0.4913 | 0.2747 |
| Multiple Logistic Regr. | 0.6706 | 0.7912 | 0.5499 | 0.3378 |
| kNN | 0.6736 | 0.8044 | 0.5428 | 0.3472 |
| Gradient Boosting | 0.7132 | 0.8329 | 0.5935 | 0.4280 |
| ANN | 0.5835 | 0.7619 | 0.4052 | 0.1729 |

(The Table S3 describes the comparative analysis of performance of different machine learning models for Dataset 1 with Randomization: Euler; K= 3)

**Table S4:** Performance Evaluation of Dataset 1 (Randomization: Linear; K= 1)

| Model | Balanced Accuracy | Specificity | Recall | MCC |
| --- | --- | --- | --- | --- |
| SVM | 0.7398 | 0.8519 | 0.6277 | 0.4806 |
| Regression Tree | 0.6596 | 0.7997 | 0.5194 | 0.3205 |
| Random Forest | 0.7412 | 0.8508 | 0.6316 | 0.4825 |
| Naive Bayes | 0.6169 | 0.7750 | 0.4588 | 0.2288 |
| Multiple Logistic Regr. | 0.7160 | 0.8388 | 0.5932 | 0.4345 |
| kNN | 0.7044 | 0.8264 | 0.5825 | 0.4086 |
| Gradient Boosting | 0.7427 | 0.8533 | 0.6321 | 0.4866 |
| ANN | 0.6244 | 0.7750 | 0.4739 | 0.2547 |

(The Table S4 describes the comparative analysis of performance of different machine learning models for Dataset 1 with Randomization: Linear; K= 1)

**Table S5:** Performance Evaluation of Dataset 1 (Randomization: Linear; K= 2)

| Model | Balanced Accuracy | Specificity | Recall | MCC |
| --- | --- | --- | --- | --- |
| SVM | 0.7164 | 0.8338 | 0.5989 | 0.4331 |
| Regression Tree | 0.6378 | 0.7736 | 0.5020 | 0.2759 |
| Random Forest | 0.7175 | 0.8330 | 0.6019 | 0.4338 |
| Naive Bayes | 0.6258 | 0.7771 | 0.4746 | 0.2477 |
| Multiple Logistic Reg. | 0.6891 | 0.8215 | 0.5567 | 0.3784 |
| kNN | 0.6675 | 0.7980 | 0.5370 | 0.3344 |
| Gradient Boosting | 0.7193 | 0.8364 | 0.6021 | 0.4389 |
| ANN | 0.6274 | 0.7723 | 0.4825 | 0.2057 |

(The Table S5 describes the comparative analysis of performance of different machine learning models for Dataset 1 with Randomization: Linear; K= 2)

**Table S6:**
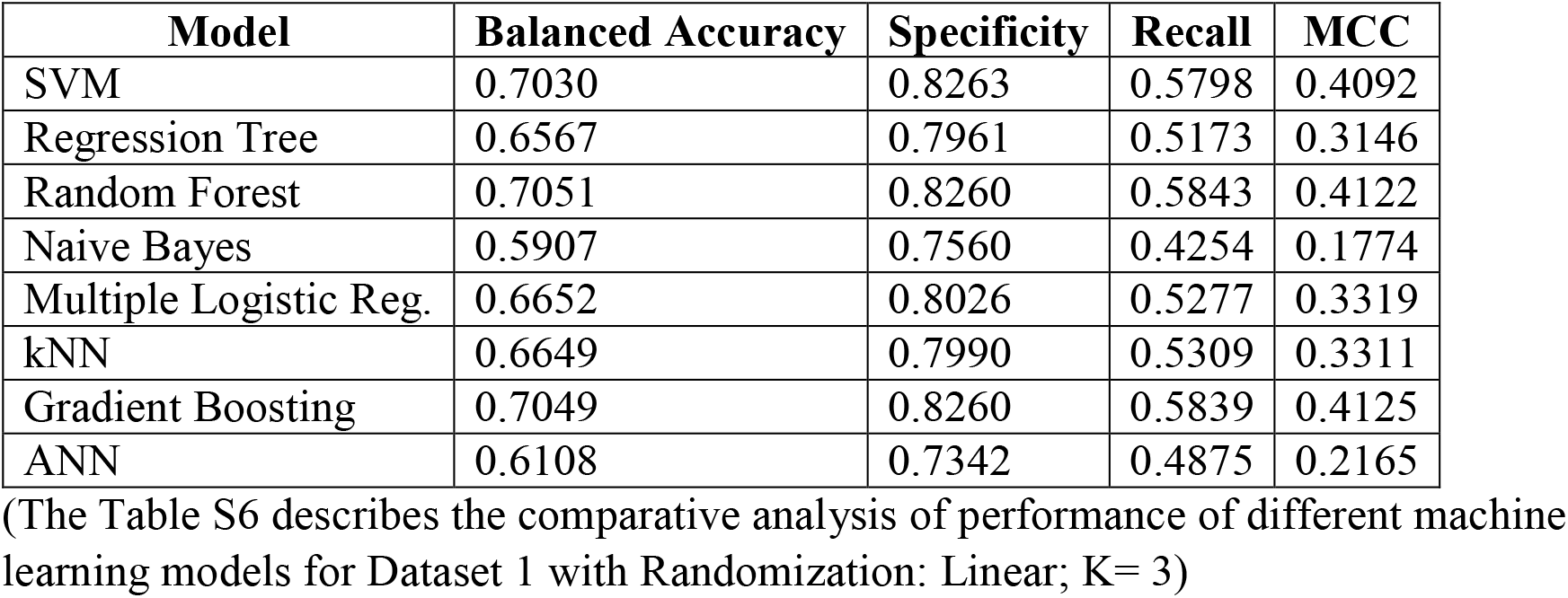
Performance Evaluation of Dataset 1 (Randomization: Linear; K= 3)

| Model | Balanced Accuracy | Specificity | Recall | MCC |
| --- | --- | --- | --- | --- |
| SVM | 0.7030 | 0.8263 | 0.5798 | 0.4092 |
| Regression Tree | 0.6567 | 0.7961 | 0.5173 | 0.3146 |
| Random Forest | 0.7051 | 0.8260 | 0.5843 | 0.4122 |
| Naive Bayes | 0.5907 | 0.7560 | 0.4254 | 0.1774 |
| Multiple Logistic Reg. | 0.6652 | 0.8026 | 0.5277 | 0.3319 |
| kNN | 0.6649 | 0.7990 | 0.5309 | 0.3311 |
| Gradient Boosting | 0.7049 | 0.8260 | 0.5839 | 0.4125 |
| ANN | 0.6108 | 0.7342 | 0.4875 | 0.2165 |

(The Table S6 describes the comparative analysis of performance of different machine learning models for Dataset 1 with Randomization: Linear; K= 3)

**Table S7:** Performance Evaluation of Dataset 1 (Randomization: Markov; K= 1)

| Model | Balanced Accuracy | Specificity | Recall | MCC |
| --- | --- | --- | --- | --- |
| SVM | 0.7203 | 0.8375 | 0.6032 | 0.4402 |
| Regression Tree | 0.6690 | 0.8019 | 0.5361 | 0.3383 |
| Random Forest | 0.7252 | 0.8391 | 0.6114 | 0.4474 |
| Naive Bayes | 0.6394 | 0.7900 | 0.4887 | 0.2739 |
| Multiple Logistic Reg. | 0.6606 | 0.7991 | 0.5222 | 0.3214 |
| kNN | 0.6741 | 0.7987 | 0.5496 | 0.3419 |
| Gradient Boosting | 0.7257 | 0.8432 | 0.6083 | 0.4515 |
| ANN | 0.6441 | 0.7667 | 0.5214 | 0.2629 |

(The Table S7 describes the comparative analysis of performance of different machine learning models for Dataset 1 with Randomization: Markov; K= 1)

**Table S8:** Performance Evaluation of Dataset 1 (Randomization: Markov; K= 2)

| Model | Balanced Accuracy | Specificity | Recall | MCC |
| --- | --- | --- | --- | --- |
| SVM | 0.7028 | 0.8219 | 0.5838 | 0.4050 |
| Regression Tree | 0.6627 | 0.7943 | 0.5312 | 0.3242 |
| Random Forest | 0.7132 | 0.8292 | 0.5972 | 0.4238 |
| Naive Bayes | 0.6309 | 0.7803 | 0.4814 | 0.2619 |
| Multiple Logistic Reg. | 0.6436 | 0.7760 | 0.5112 | 0.2878 |
| kNN | 0.6611 | 0.7837 | 0.5385 | 0.3169 |
| Gradient Boosting | 0.7096 | 0.8279 | 0.5912 | 0.4186 |
| ANN | 0.6440 | 0.7588 | 0.5292 | 0.2785 |

(The Table S8 describes the comparative analysis of performance of different machine learning models for Dataset 1 with Randomization: Markov; K= 2)

**Table S9:**
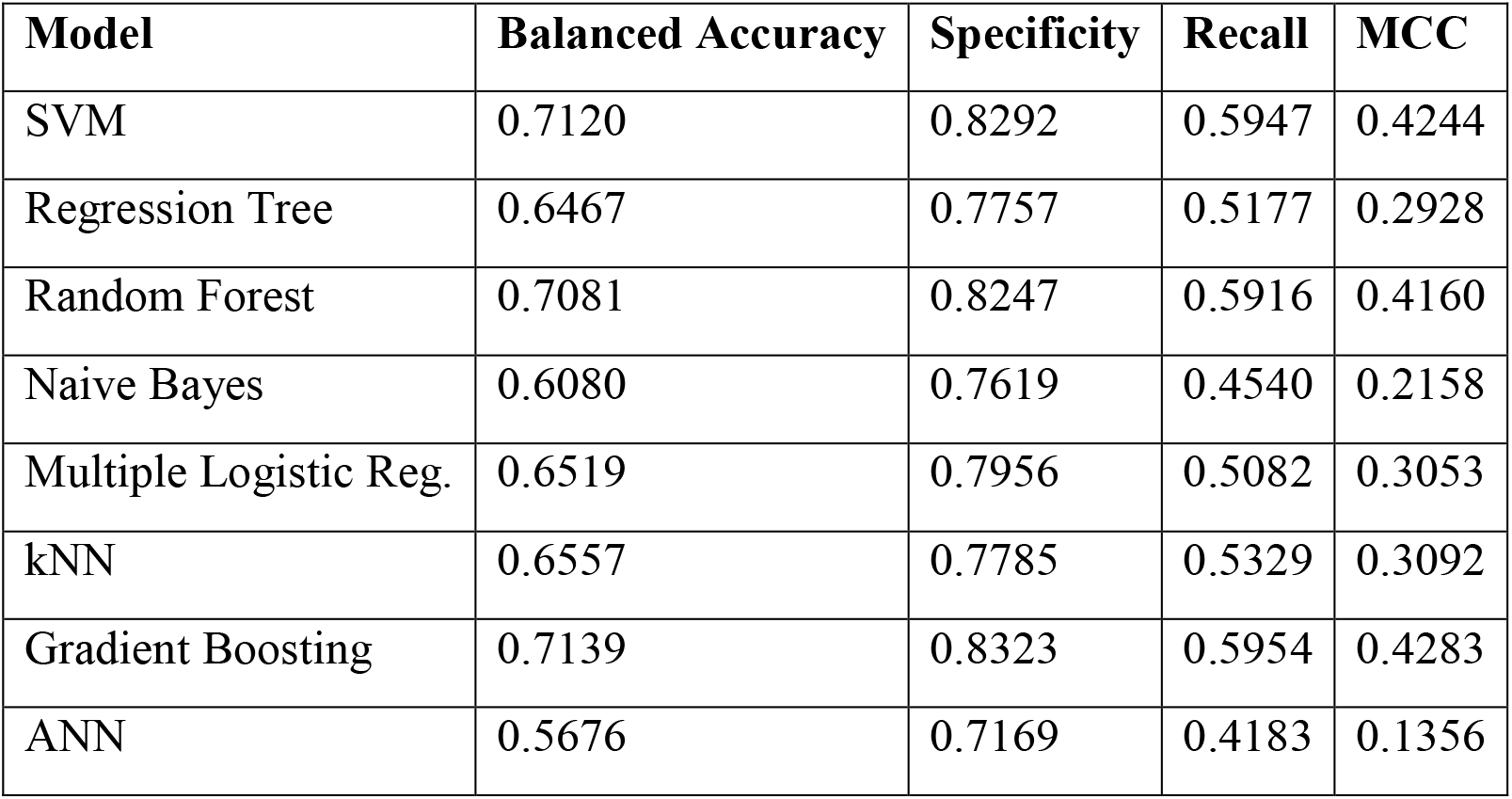
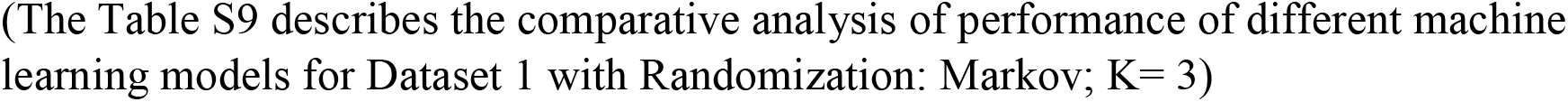
Performance Evaluation of Dataset 1 (Randomization: Markov; K= 3)

| Model | Balanced Accuracy | Specificity | Recall | MCC |
| --- | --- | --- | --- | --- |
| SVM | 0.7120 | 0.8292 | 0.5947 | 0.4244 |
| Regression Tree | 0.6467 | 0.7757 | 0.5177 | 0.2928 |
| Random Forest | 0.7081 | 0.8247 | 0.5916 | 0.4160 |
| Naive Bayes | 0.6080 | 0.7619 | 0.4540 | 0.2158 |
| Multiple Logistic Reg. | 0.6519 | 0.7956 | 0.5082 | 0.3053 |
| kNN | 0.6557 | 0.7785 | 0.5329 | 0.3092 |
| Gradient Boosting | 0.7139 | 0.8323 | 0.5954 | 0.4283 |
| ANN | 0.5676 | 0.7169 | 0.4183 | 0.1356 |

(The Table S9 describes the comparative analysis of performance of different machine learning models for Dataset 1 with Randomization: Markov; K= 3)

**Table S10:**
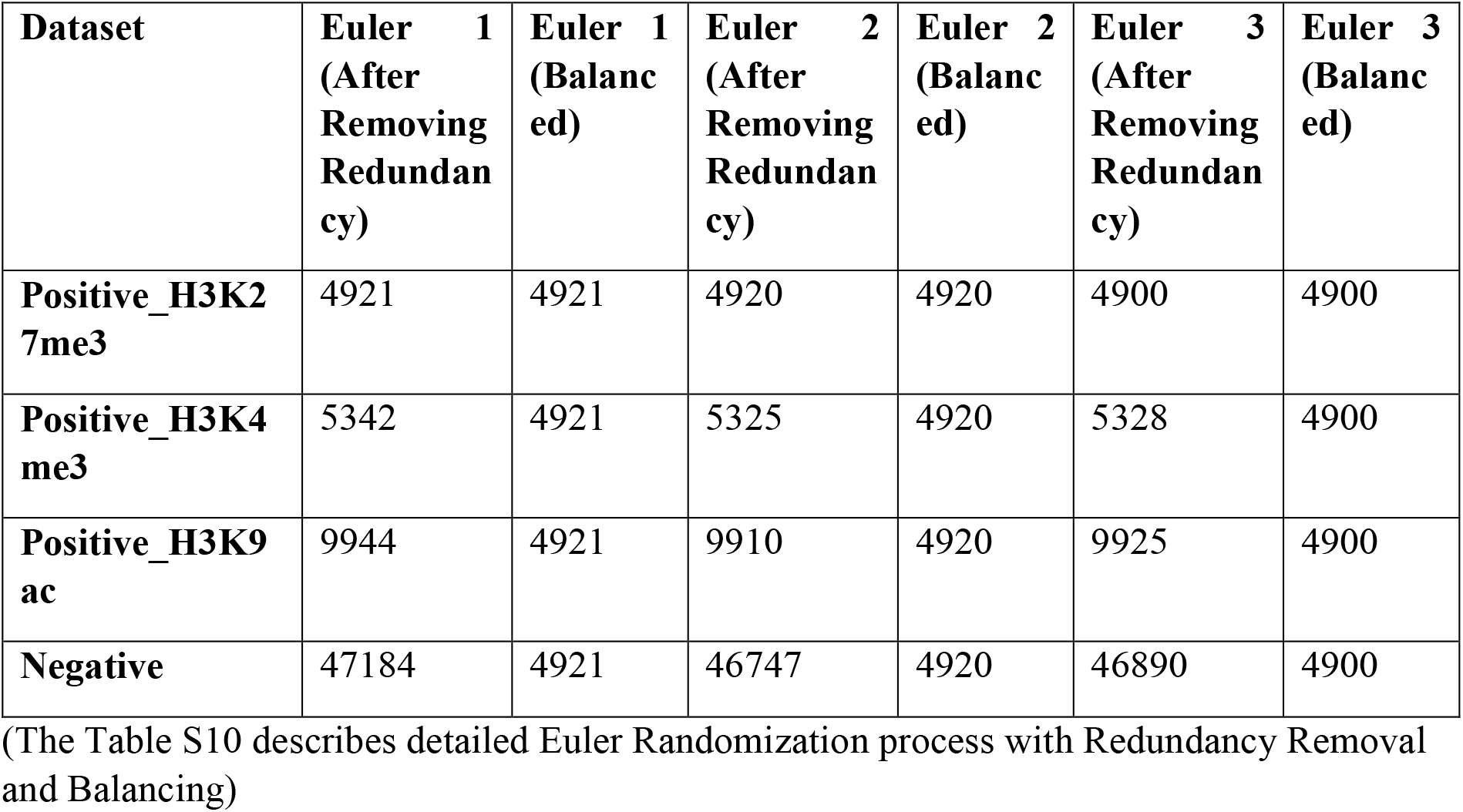
Euler Randomization.

| **Dataset** | **Euler 1**<br>**(After**<br>**Removing**<br>**Redundan**<br>**cy)** | **Euler 1**<br>**(Balanc**<br>**ed)** | **Euler 2**<br>**(After**<br>**Removing**<br>**Redundan**<br>**cy)** | **Euler 2**<br>**(Balanc**<br>**ed)** | **Euler 3**<br>**(After**<br>**Removing**<br>**Redundan**<br>**cy)** | **Euler 3**<br>**(Balanc**<br>**ed)** |
| --- | --- | --- | --- | --- | --- | --- |
| **Positive_H3K27me3** | 4921 | 4921 | 4920 | 4920 | 4900 | 4900 |
| **Positive_H3K4me3** | 5342 | 4921 | 5325 | 4920 | 5328 | 4900 |
| **Positive_H3K9ac** | 9944 | 4921 | 9910 | 4920 | 9925 | 4900 |
| **Negative** | 47184 | 4921 | 46747 | 4920 | 46890 | 4900 |

(The Table S10 describes detailed Euler Randomization process with Redundancy Removal and Balancing)

**Table S11:** Linear Randomization.

| **Dataset** | **Linear 1**<br>**(After**<br>**Removing**<br>**Redundan**<br>**cy)** | **Linear 1**<br>**(Balanc**<br>**ed)** | **Linear 2**<br>**(After**<br>**Removing**<br>**Redundan**<br>**cy)** | **Linear 2**<br>**(Balanc**<br>**ed)** | **Linear 3**<br>**(After**<br>**Removing**<br>**Redundan**<br>**cy)** | **Linear 3**<br>**(Balanc**<br>**ed)** |
| --- | --- | --- | --- | --- | --- | --- |
| **Positive_H3K27me3** | 4931 | 4931 | 4938 | 4938 | 4902 | 4902 |
| **Positive_H3K4me3** | 5341 | 4931 | 5335 | 4938 | 5329 | 4902 |
| **Positive_H3K9ac** | 9950 | 4931 | 9943 | 4938 | 9922 | 4902 |
| **Negative** | 47119 | 4931 | 47072 | 4938 | 46913 | 4902 |

(The Table S11 describes detailed Linear Randomization process with Redundancy Removal and Balancing)

**Table S12:** Markov Randomization.

| **Dataset** | **Markov 1**<br>**(After** | **Markov**<br>**1** | **Markov 2**<br>**(After** | **Markov**<br>**2** | **Markov 3**<br>**(After** | **Markov**<br>**3** |
| --- | --- | --- | --- | --- | --- | --- |

|  | **Removing**<br>**Redundan**<br>**cy)** | **(Balanc**<br>**ed)** | **Removing**<br>**Redundan**<br>**cy)** | **(Balanc**<br>**ed)** | **Removing**<br>**Redundan**<br>**cy)** | **(Balanc**<br>**ed)** |
| --- | --- | --- | --- | --- | --- | --- |
| **Positive_H3K2**<br>**7me3** | 4935 | 4935 | 4935 | 4935 | 4938 | 4938 |
| **Positive_H3K4**<br>**me3** | 5336 | 4935 | 5349 | 4935 | 5341 | 4938 |
| **Positive_H3K9**<br>**ac** | 9951 | 4935 | 9946 | 4935 | 9950 | 4938 |
| **Negative** | 47207 | 4935 | 46540 | 4935 | 46646 | 4938 |

(The Table S12 describes detailed Markov Randomization process with Redundancy Removal and Balancing)

**Figure S1:**
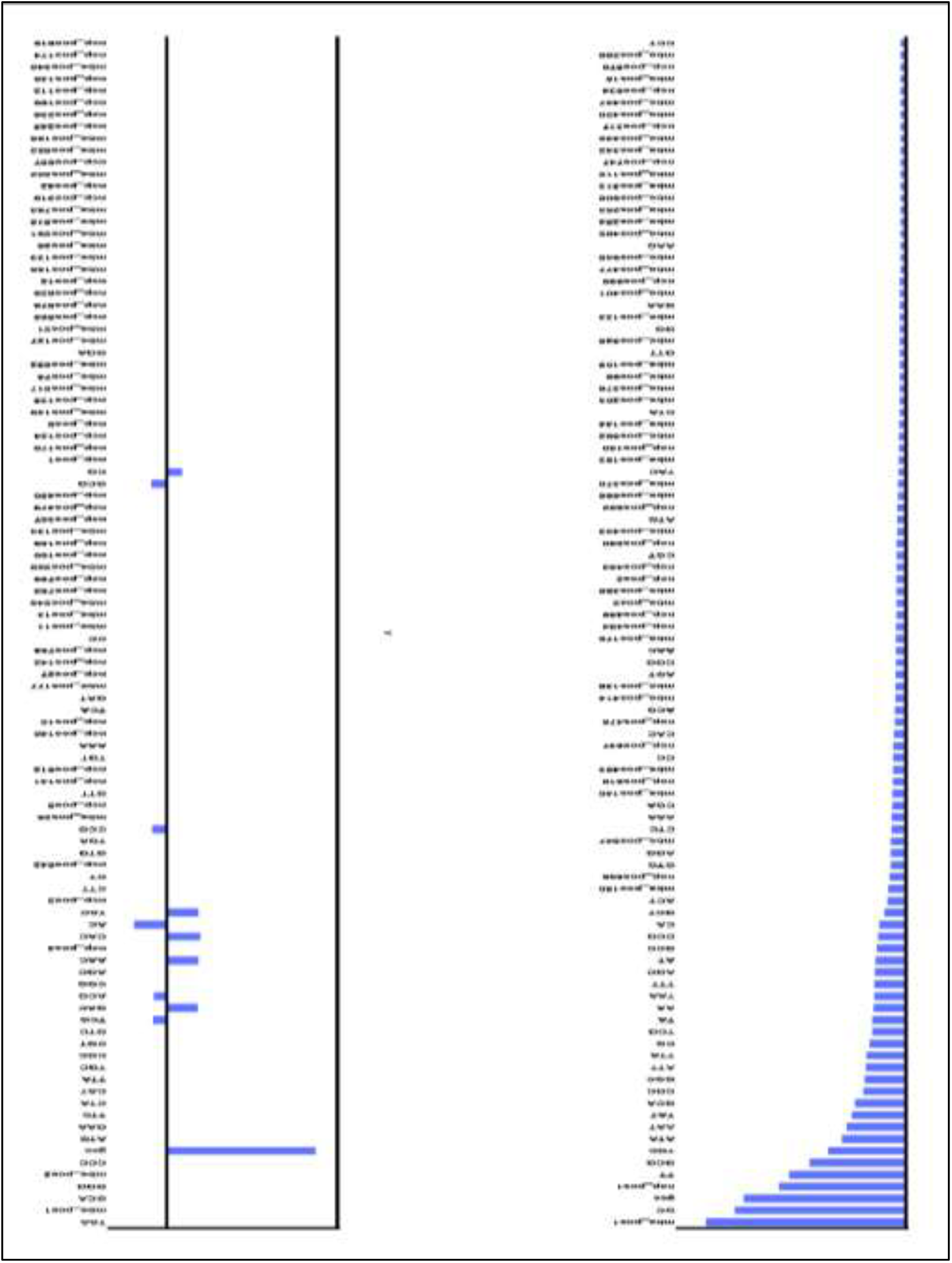
Feature Importance Score obtained from RF and SwR. (The Figure S1 describes comparative graphical representation of feature importance score for Random Forest and Stepwise Regression based feature selection criterion)

